# HNI9 and HY5 maintain ROS homeostasis under high nitrogen provision in Arabidopsis

**DOI:** 10.1101/479030

**Authors:** Fanny Bellegarde, Amel Maghiaoui, Jossia Boucherez, Gabriel Krouk, Laurence Lejay, Liên Bach, Alain Gojon, Antoine Martin

## Abstract

**One sentence summary:** Excessive N nutrition leads to ROS accumulation, and requires the function of major transcriptional regulators to maintain plants under physiological conditions.

**Author contributions:** An.M. and A.G. conceived research plans and supervised the experiments; F.B, Am.M., J.B., L.L., L.B. and An.M performed most of the experiments; F.B, Am.M., J.B., G.K., L.L., L.B. and An.M analyzed the data; An.M. wrote the article with contributions of all the authors.

**Competing interests:** The authors declare no competing financial interests.

**Summary:** Reactive Oxygen Species (ROS) can accumulate in cells at excessive levels, leading to unbalanced redox status and to a potential oxidative stress, which can have damaging effects to the molecular components of plant cells. Several environmental conditions have been described as causing an elevation of ROS production in plants. Consequently, this requires the expression of detoxification responses in order to maintain ROS homeostasis at physiological levels. In case of mis-regulation of the detoxification systems, oxidative stress can lead ultimately to growth retardation and developmental defects. Here, we demonstrate that Arabidopsis plants growing under high nitrogen environment have to express a set of genes involved in detoxification of ROS in order to maintain ROS at physiological levels. We show that the chromatin factor HNI9 is an important actor of this response, required for the expression of these detoxification genes. Mutation in HNI9 leads to elevated ROS levels, and to ROS-dependent phenotypic defects under high but not low N provision. In addition, we identify HY5 as one of the major transcription factors also required for the expression of this detoxification program under high N condition. Our results demonstrate the requirement of a balance between N nutrition and ROS production, and identified the first major regulators required to control ROS homeostasis under excessive N nutrition.

## Introduction

Reactive Oxygen Species (ROS) are integral part of plant metabolism, generated as by-products of a large range of enzymatic reactions. The dynamics of ROS in plant cells corresponds to two distinct scenarios. First, ROS are involved in numerous signaling pathways, and thus their dynamic affects a lot of developmental and physiological processes, like cell differentiation or response to biotic and abiotic stresses (Noctor et al., 2018). In such cases, variations in ROS are generally dynamic and transient, and occur at moderate concentration. However, on another hand, ROS can also accumulate in cells at excessive and constant levels, leading to unbalanced redox status and to potential oxidative stress (Schieber and Chandel, 2014). At the cellular level, excessive accumulation of ROS can lead to oxidation and to damage of many essential molecules (Moller et al., 2007). For instance, oxidation of enzymatic proteins often leads to loss of activity, oxidation of DNA can lead to degradation or mutations, and oxidation of lipids leads to disorganization of cellular membranes. The maintenance of ROS homeostasis is therefore essential to keep cell integrity. ROS levels are also associated with a range of physiological and developmental phenotypes. For instance, functioning of the shoot apical meristem is largely influenced by the redox status (Schippers et al., 2016), and many developmental processes imply interactions between ROS and phytohormones (Considine and Foyer, 2014). Several reports have also demonstrated a role for ROS in root growth and development (Tsukagoshi, 2016). As a consequence, mutations leading to excessive ROS production or exogenous application of ROS at high concentration lead to retardation of root growth in Arabidopsis (Dunand et al., 2007; Tsukagoshi et al., 2010; Zhao et al., 2016). Altogether, these observations highlight that controlling the level of ROS is crucial for plant growth, development and physiology.

Given the importance of ROS homeostasis, plants possess a large range of mechanisms that are able to remove ROS from the cells. A main part of this antioxidant response corresponds to enzymes, such as peroxidases, that use reducing power in oxidation/reduction reactions to decrease the cell redox status (Moller et al., 2007). These enzymes work in complex systems also involving essential antioxidant molecules, like ascorbic acid, glutathione or thiamin, that are known to protect against oxidative stress (Tunc-Ozdemir et al., 2009; Ramírez et al., 2013). At the molecular level, transcriptional induction of ROS-sensitive genes in plants constitutes a major part of the response to ROS overproduction (Willems et al., 2016). However, in agreement with the high number of genes coding for components of antioxidant responses in Arabidopsis, a specificity of response seems to occur in function of the signals at the origin of ROS production. Indeed, signaling pathways and genes involved in ROS homeostasis in response to stress largely vary according to the type of stress encountered by the plant (Apel and Hirt, 2004). However, many of the actors that drive each transcriptional response and its specificity remain to be identified.

ROS production has been observed in plants in response to many environmental factors, with both biotic interactions and abiotic stress often leading to ROS signaling (Baxter et al., 2014). Concerning abiotic constraints, drought, salinity or heat stress are known to induce the production of ROS (Choudhury et al., 2017). However, these responses correspond to cases where ROS contribute to a signaling pathways, but do not accumulate and generate oxidative stress. In opposition, several examples of abiotic stress have been directly linked to an elevation of ROS, which is eventually perceived as a perturbation of the plant redox balance. This is the case for nutrients deprivation, where potassium, nitrogen (N) or phosphorus starvation rapidly leads to an accumulation of H_2_O_2_ in the roots (Shin and Schachtman, 2004; Shin et al., 2005). This suggest that the redox status of plants can be very sensitive to changes in the nutritional environment.

Previously, the chromatin factor HNI9 has been shown to be involved in the response to high N provision in Arabidopsis. Mutations in *HNI9* leads to an increase in transcripts level corresponding to the nitrate transporter *NRT2.1* under high N provision, where this gene is normally strongly downregulated (Widiez et al., 2011). However, the direct relationship between HNI9 and *NRT2.1* has not been demonstrated. Furthermore, several reports in plants, animals and yeast demonstrated that HNI9 is a positive regulator of gene expression (Yoh et al., 2008; Li et al., 2010; Chen et al., 2012; Wang et al., 2013; Wang et al., 2014), which is not consistent with the hypothesis of a direct role of HNI9 on *NRT2.1* downregulation. Here, we reexamined the function of HNI9 in the response of plants to high N provision, and we show an unexpected role of HNI9 in the regulation of ROS homeostasis under high N provision, through the induction of a subset of genes involved in detoxification. We also identify the transcription factor HY5 as an important component of this response, and highlight a new interaction between plant nutrition and ROS homeostasis.

## Results

### HNI9 is required to upregulate the expression of a set of genes involved in redox processes in response to high N provision

In order to reexamine the role of HNI9 in the response to high N provision, we compared transcriptomic data of WT and *hni9-1* mutant plants under low nitrate and high N provisions (Widiez et al., 2011). As HNI9 is a positive regulator of gene expression, we selected genes that are induced by high N provision in WT, and looked for those that do not display such induction in the *hni9-1* mutant line. Using these criteria, we obtained a list of 108 genes induced under high N condition in a HNI9-dependent manner (Table S1). In order to determine which biological functions are affected by the HNI9 mutation under high N provision, we performed a gene ontology analysis using this list of 108 genes. Among the biological functions overrepresented, we found that several categories related to redox processes, including oxidoreductase activity, peroxidase activity, or antioxidant activity, were the most significantly represented (Figure 1, Table 1). In addition, using further manual curation, we observed that many other genes present in other GO categories might also encode for enzymes or proteins associated with ROS detoxification or with antioxidant biosynthesis pathways (Table S1). Therefore, we hypothesized that high N provision could challenge ROS homeostasis in plants, and that HNI9 could be required in order to induce a set of genes involved in cellular detoxification.

**Figure 1:**
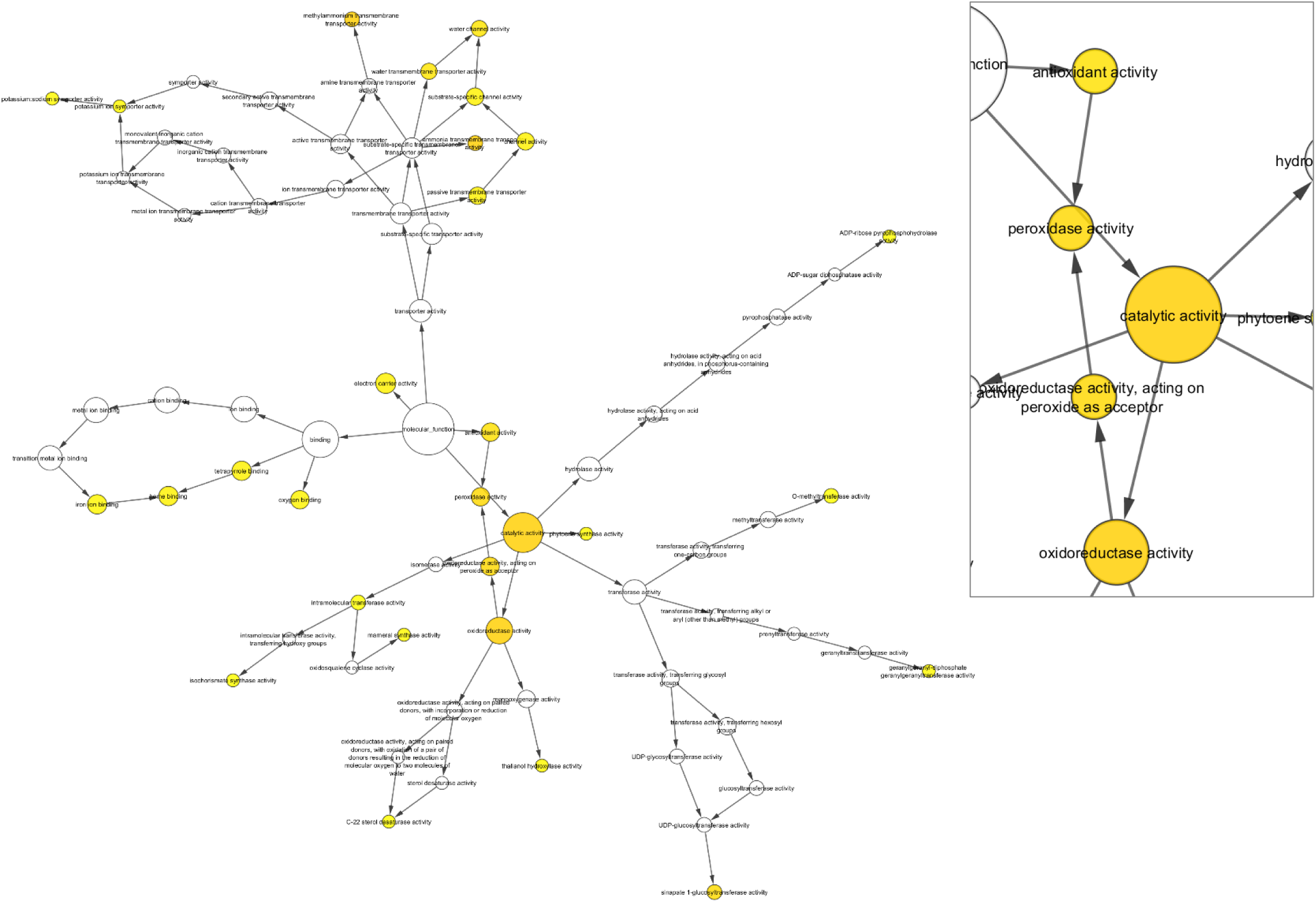
Genes induced by high N provision in HNI9 dependent manner are associated with redox processes. Functional network realized from the list of 108 genes induced specifically under high N condition (10 mM NH_4_NO_3_) in a HNI9 dependent manner. Orange and yellow circles correspond to p-values lower than 0.01 and 0.05, respectively (see also Table 1 for details). Insert in the right corresponds to the central hub of the network with biological function associated with antioxidant activities.

**Table 1:**
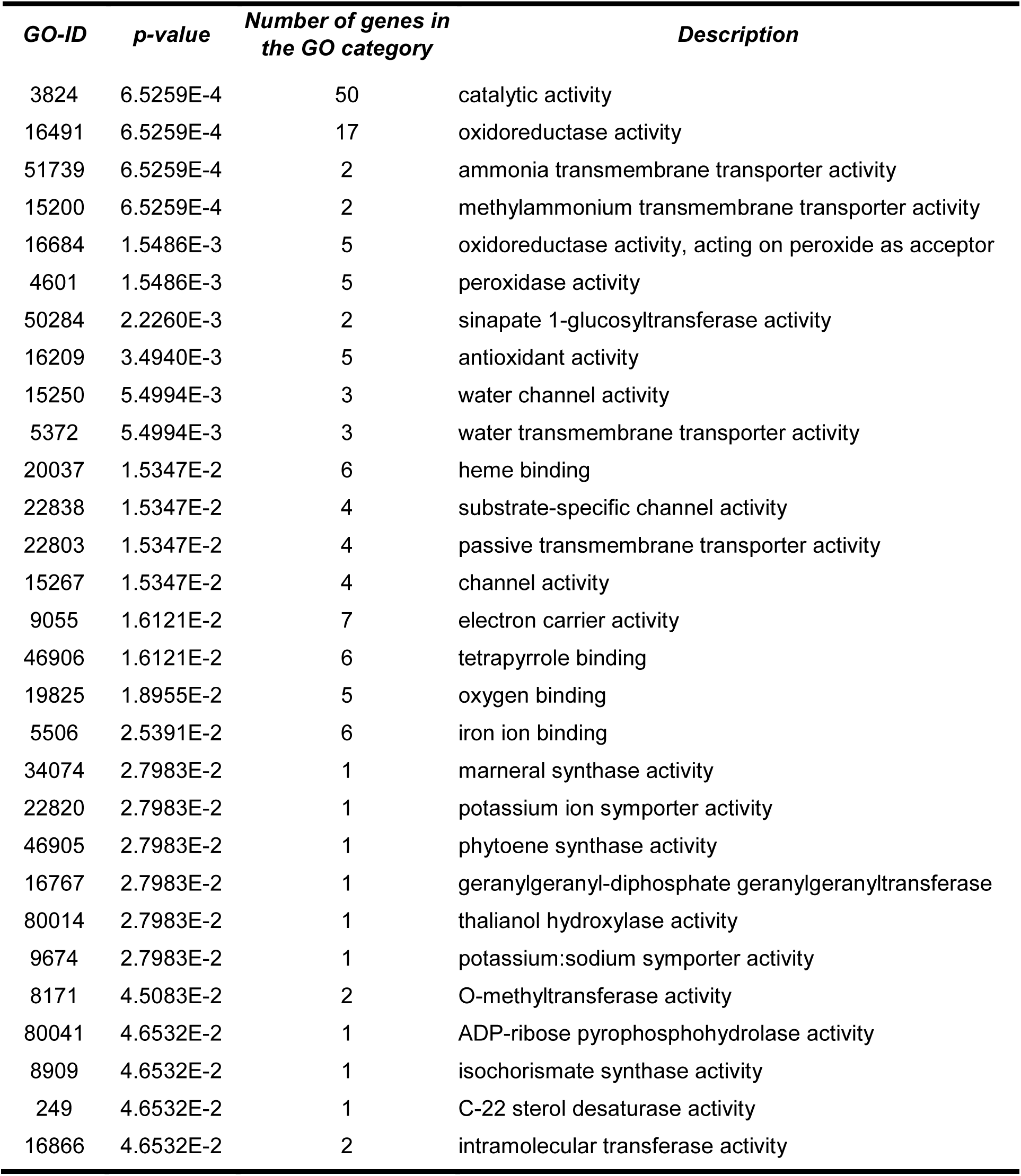
Gene ontology enrichment identified in HNI9-dependent genes induced by high N conditions (10 mM NH_4_NO_3_). The gene ontology term enrichment analysis was performed under Cytoscape environment by BINGO software using the Molecular Function ontology file. Details of the analysis are provided in Supplementary File 1.

### HNI9 mutation leads to higher ROS levels and to ROS-dependent phenotypes under high N provision

In order to assess the role of HNI9 in ROS homeostasis under high N provision, we measured H_2_O_2_ and total ROS levels in WT and *hni9-1* mutant lines under low nitrate and high N provisions. In WT line, H_2_O_2_ levels remained constant whatever N provision, whereas total ROS levels slightly increase under high N provision, yet not significantly, showing that ROS homeostasis is maintained from low to high N provision (Figure 2A, B). In contrast, ROS accumulation displayed significant changes in the *hni9-1* mutant line. The levels of H_2_O_2_ and total ROS in the *hni9-1* mutant line were identical to that of WT under low N provision, but they increased significantly under high N provision (Figure 2A, B). To confirm these results, we used an Arabidopsis line expressing the *GRX1-roGFP2* construct (hereafter mentioned as *roGFP2*), in order to visualize ROS levels *in planta*. RoGFP2 is a redox-sensitive probe, which allows to monitor plant redox status and in particular to address H_2_O_2_ levels *in vivo* (Meyer et al., 2007; Marty et al., 2009). In WT plants under high N provision, fluorescence signal of roGFP2 was very low, in agreement with our measurements of H_2_O_2_ and ROS levels (Figure 2C). In contrast, fluorescence signal of roGFP2 in *hni9-1* plants was strong, corresponding to the elevation of H_2_O_2_ and ROS levels measured in this line (Figure 2D). Quantification of roGFP2 signals in each lines validates a strongly significant increase in ROS levels in *hni9-1* mutant line as compared to the WT (Figure S1). Altogether, these results led us to conclude that HNI9 contributes to prevent ROS overaccumulation under high N supply, likely through the induction of a set of genes required to detoxify ROS produced under high N condition.

**Figure 2:**
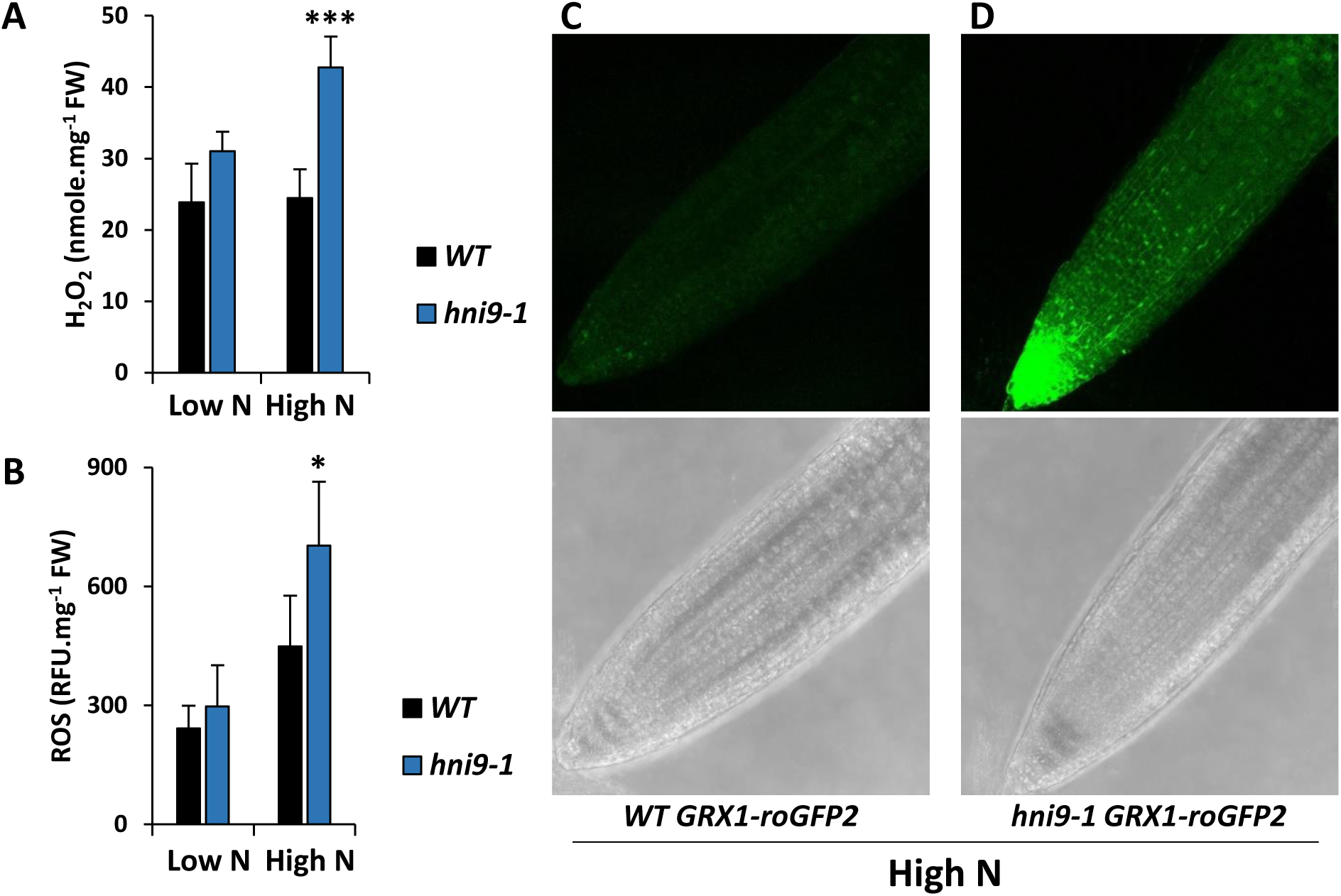
ROS levels are higher in *hni9-1* under high N provision. Measurement of H_2_O_2_(A) and ROS (B) in roots of WT and *hni9-1* lines under low (0.3 mM NO_3_-) and high N (10 mM NH_4_NO_3_) provision. Statistical significance was computed using a two-tailed Student’s t-test. Significance cutoff: *p < 0.05, **p < 0.01, ***p < 0.001. Visualization of H_2_O_2_ levels *in vivo* using *GRX1-roGFP2* probe in WT (C) or *hni9-1* (D) lines under high N conditions.

We next looked whether upregulation of *NRT2.1* expression under high N supply, which is a main phenotype described for the *hni9-1* mutant line, could be linked with ROS overaccumulation in this mutant. Therefore, we investigated the expression of *NRT2.1* under two conditions associated with elevated ROS levels. First, we measured *NRT2.1* transcript levels in WT plants treated with menadione, a redox-active quinone which causes an elevation of ROS in plant roots (Lehmann et al., 2009). Our results showed that this treatment indeed results in an increase in *NRT2.1* transcript levels (Figure 3A). Then we measured *NRT2.1* transcript levels in the ascorbate-deficient mutant *vtc2*, which is affected in ROS detoxification processes and displays significantly higher H_2_O_2_ levels than in WT plants (Kotchoni et al., 2009). In comparison to WT, *NRT2.1* transcripts levels were significantly higher in the *vtc2* mutant line, as we observed for the *hni9-1* mutant (Figure 3B). These observations suggest that *NRT2.1* is induced by ROS, and thus lend support to the hypothesis that the previously reported *hni9-1* phenotype under high N condition may be due to elevation of ROS levels in this mutant. Conversely, we investigated whether well-known phenotypes associated with ROS overaccumulation were also observed in *hni9-1* plants under high N supply. In particular, elevated ROS levels have been shown to alter root growth, and thus plants growing under elevated ROS levels display a reduced primary root length (Dunand et al., 2007; Tsukagoshi et al., 2010). Therefore, we asked whether elevated ROS levels in *hni9-1* mutant line under high N provision are correlated with an alteration of root growth. We used the *vtc2* mutant line, affected in ROS detoxification processes, as a positive control to confirm that elevated ROS levels lead to a reduction of root growth under our experimental conditions. Indeed, the *vtc2* mutant line showed a reduction of primary root length as compared to the WT, regardless of the level of N provision; however, the reduction of root growth was significantly higher under high N than under low N (Figure 4). The *hni9-1* mutant line also displays a reduction of root growth in comparison to the WT (Figure 4), which is consistent with the general role of HNI9 on plant growth and development that has been previously described (Li et al., 2010). However, the reduction of root growth was also significantly higher under high N condition than under low N condition (Figure 4). Again, this suggests that phenotypic changes due to HNI9 mutation under high N supply may be explained by ROS overaccumulation. Finally, we tested whether external application of antioxidant molecule could complement the *hni9-1* root growth phenotype. We therefore measured root growth of WT and *hni9-1* lines under high N conditions, supplemented or not with thiamin, an antioxidant known to mitigate growth retardation caused by oxidative stress (Tunc-Ozdemir et al., 2009). Indeed, we observed that primary root growth defect of *hni9-1* was fully abolished in the presence of thiamin (Figure 5). This demonstrates that the presence of antioxidant can complement *hni9-1* root growth phenotype, further indicating that HNI9 affects root development under high N supply specifically through altered ROS homeostasis.

**Figure 3:**
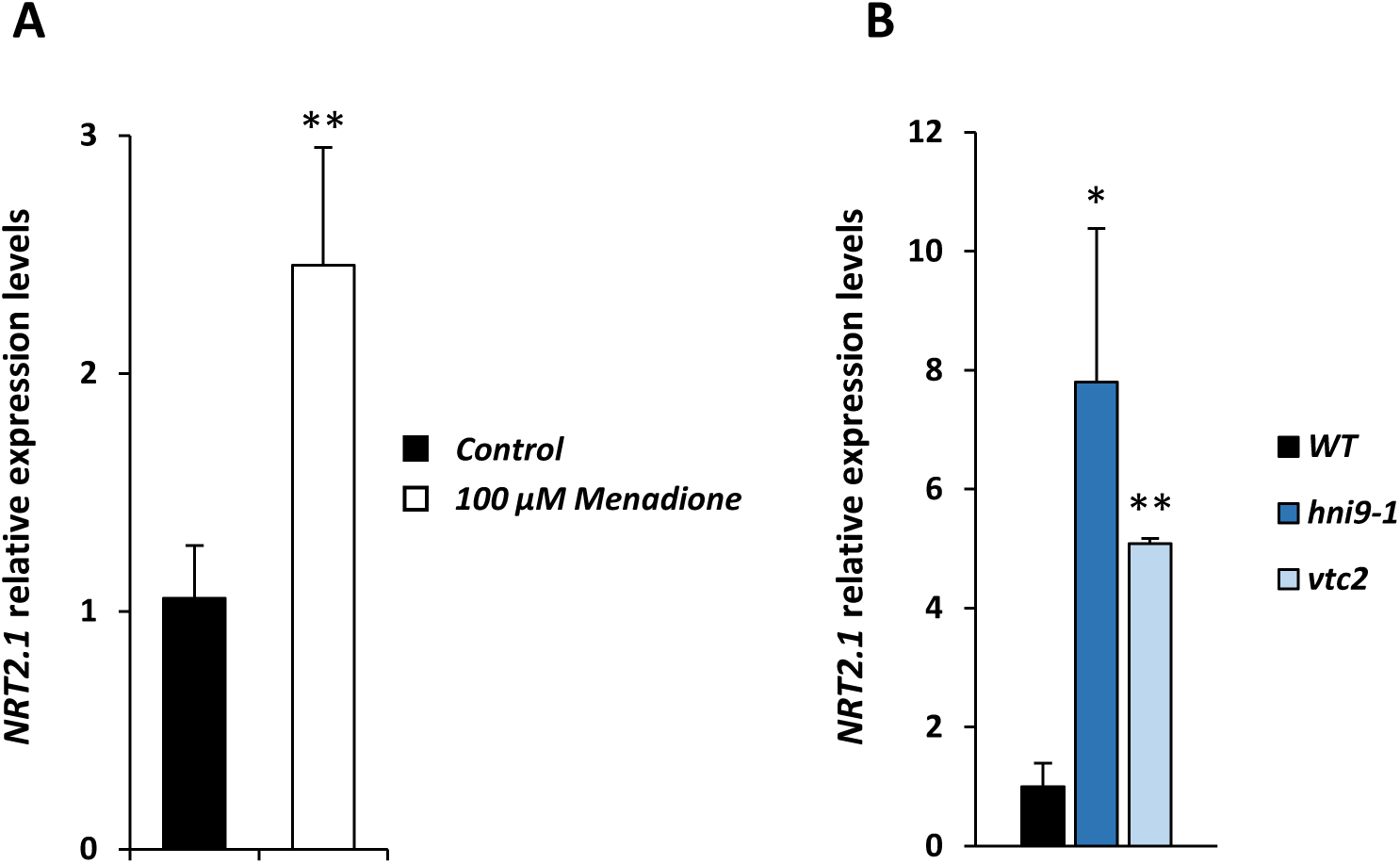
*NRT2.1* expression is sensitive to ROS homeostasis. A. Relative expression of *NRT2.1* in the presence of 100 µM menadione in roots of WT plants. Plants were grown under MS/2 medium containing 1 mM NO_3_-, and transferred on the same medium with or without menadione for 4 hours. B. Relative expression of *NRT2.1* in roots of WT, *hni9-1* and *vtc2* mutants. Plants were grown under MS/2 medium containing high N provision (10 mM NH_4_NO_3_). Statistical significance was computed using a two-tailed Student’s t-test. Significance cutoff: *p < 0.05, **p < 0.01, ***p < 0.001.

**Figure 4:**
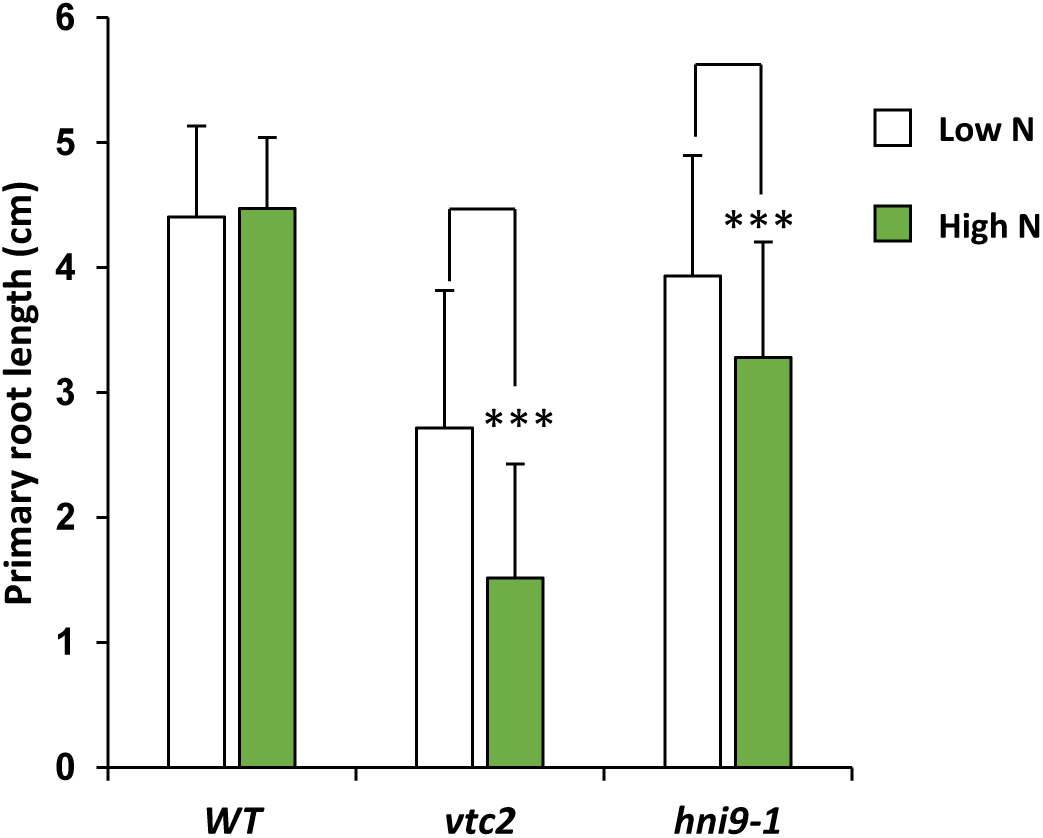
Root growth retardation is more pronounced under high N provision in *hni9-1* and *vtc2* mutants. Primary root length measurement of 7 days-old plants grown under low (0.3 mM NO_3_-) and high N (10 mM NH_4_NO_3_) provision, in WT, *hni9-1* and *vtc2* lines. The extent of root growth reduction is enhanced under high N condition, correlated with the presence of ROS in the mutant lines. Statistical significance was computed using a two-tailed Student’s t-test. Significance cutoff: *p < 0.05, **p < 0.01, ***p < 0.001.

**Figure 5:**
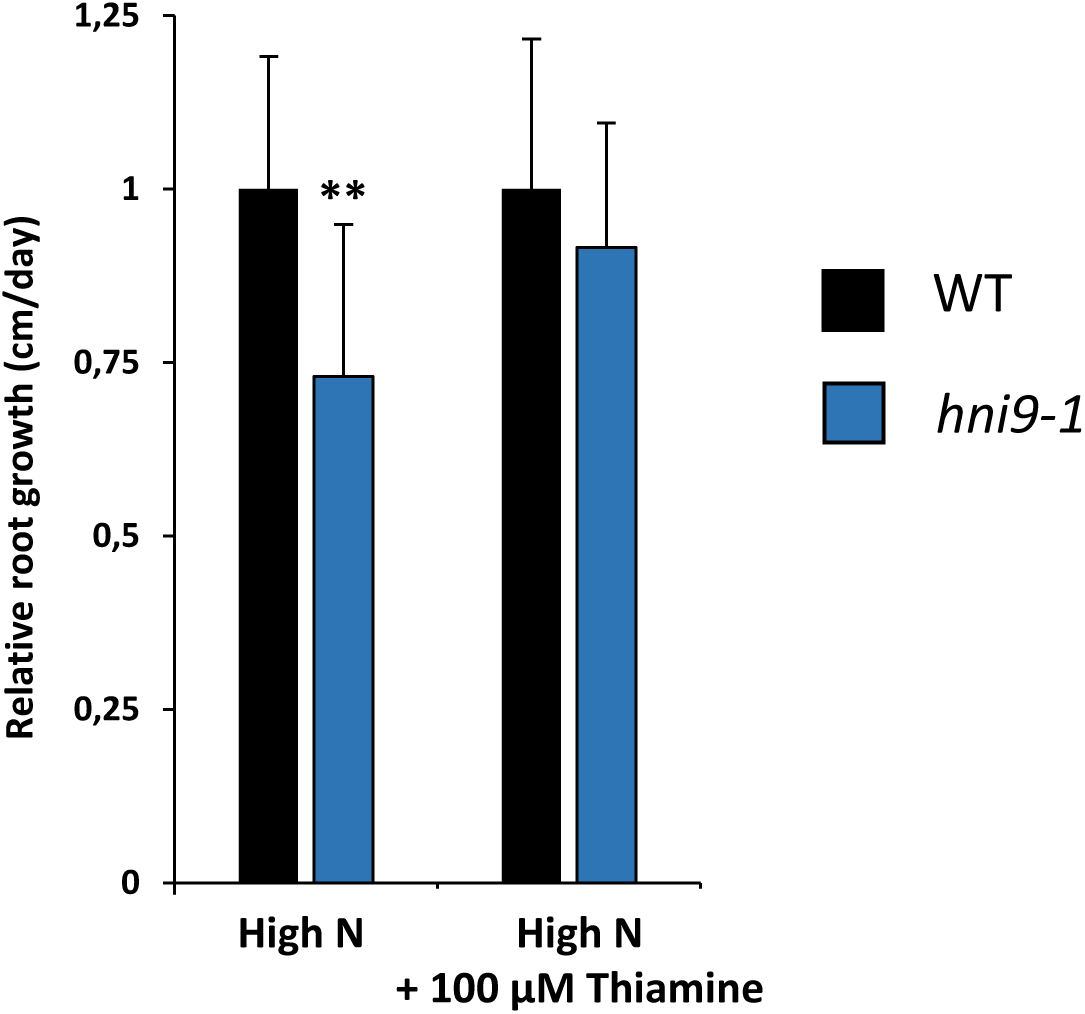
External application of antioxidant molecule complement the *hni9-1* root growth phenotype. Plants were grown under high N condition (10 mM NH_4_NO_3_) for 4 days, and transferred to the same medium with or without 100 µM thiamin. Primary root growth of WT and *hni9-1* lines was measured after 2 days of growth. Statistical significance was computed using a two-tailed Student’s t-test. Significance cutoff: *p < 0.05, **p < 0.01, ***p < 0.001.

### HNI9 is required to achieve full levels of H3K4me3 at the locus of detoxification genes

HNI9 is part of a large protein complex, including elongation and chromatin remodeling factors, which affects the histone modification state of genes (Yoh et al., 2008; Li et al., 2010). Indeed, several reports in plants and animals demonstrated that HNI9 operates at the switch between transcriptional repression and activation, in cooperation with H3K27me3 demethylases, and H3K36 methyl-transferases (Li et al., 2013; Wang et al., 2014). Therefore, we tested the hypothesis that HNI9 controls the expression of ROS detoxification genes by modulating their chromatin state in response to high N condition. To do so, we assayed the level of 3 active chromatin marks (H3K4me3, H3K9ac and H3K36me3) and 1 repressive chromatin mark (H3K27me3) at the locus of several HNI9-dependent genes involved in ROS detoxification in response to high N condition. Here, the results showed that the level of H3K27me3 and H3K36me3 were not changed in the *hni9-1* mutant at the locus of HNI9-dependent detoxification genes (Figure 6). In addition, the levels of H3K9ac, associated with active transcription were also generally similar between WT and *hni9-1*. However, we observed that H3K4me3 levels were globally lower in the *hni9-1* mutant at the locus of HNI9-dependent detoxification genes (Figure 6). We therefore conclude that HNI9 may induce the expression of ROS detoxification genes by regulating the profile of H3K4me3.

**Figure 6:**
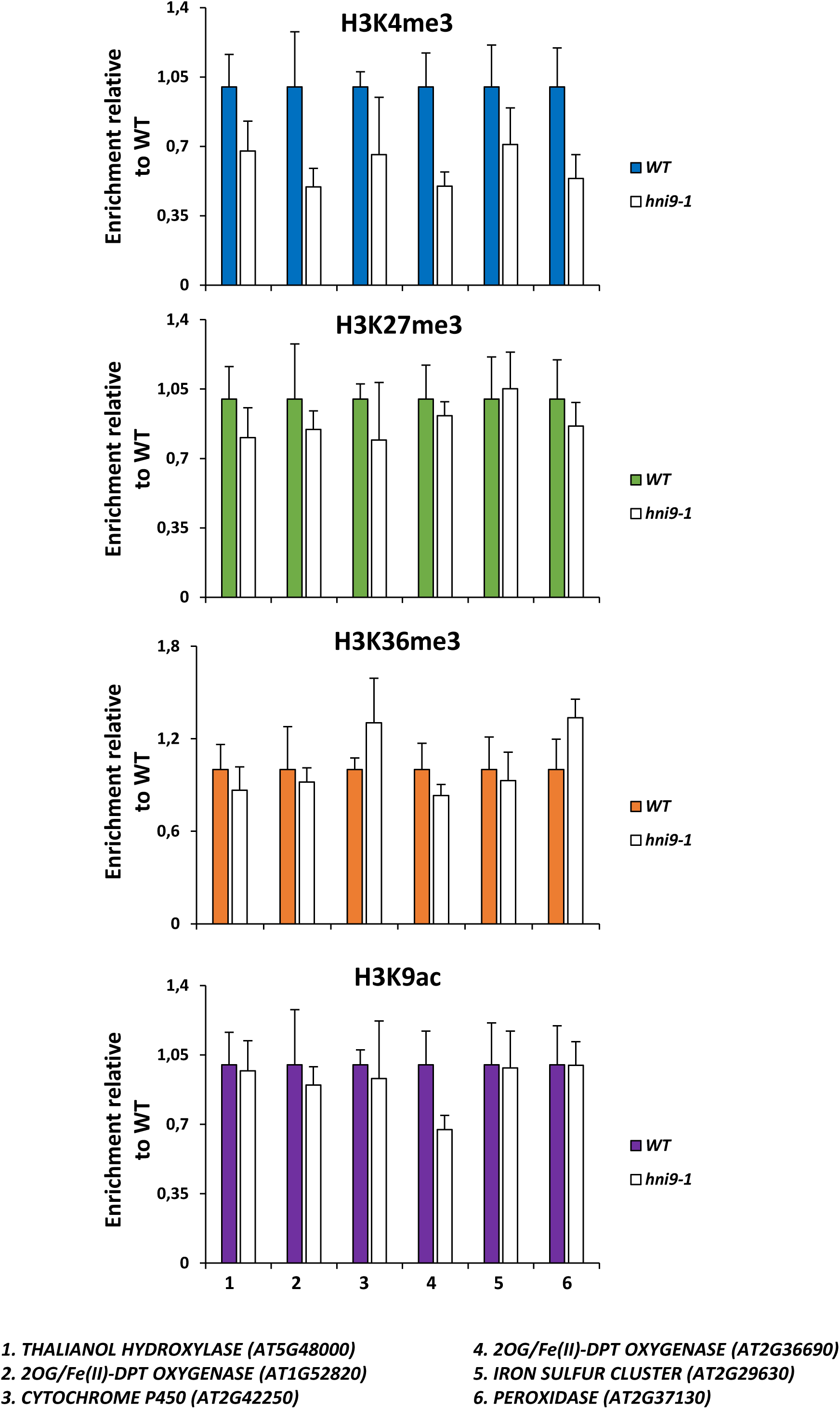
Mutation in HNI9 is associated with reduction of H3K4me3 at the loci of detoxification genes. ChIP analysis of H3K4me3, H3K27me3, H3K36me3 and H3K9ac in *WT* and *hni9-1* roots of 7 days-old plants grown under high N provision (10 mM NH_4_NO_3_). Quantification by qRT-PCR is shown as the percentage of H3K4me3, H3K36me3, H3K27me3 or H3K9ac over H3 at target loci, normalized by the percentage of H3K4me3, H3K36me3, H3K27me3 or H3K9ac over H3 at positive controls (*ACT2* for H3K4me3, H3K36me3, and H3K9ac, *LEC2* for H3K27me3). Data are presented relative to the WT level. Error bars represent standard errors of the mean based on at least 3 biological replicates.

### Analysis of HNI9-dependent genes involved in ROS detoxification under high N condition reveals the role of HY5 in the control of ROS homeostasis

HNI9 is a generic regulator of gene expression, as it does not provide by itself any sequence specificity to its target loci. Specificity of response thus requires the action of transcription factors, to drive chromatin remodeling complexes to target loci (Li et al., 2010). Therefore, we tested whether promoter sequence analysis of genes involved in ROS detoxification under high N could reveal the implication of transcription factors. We submitted the promoter sequences from the 108 genes induced by high N and dependent on HNI9 to MEME software in order to find putative conserved cis regulatory elements (CREs). This analysis reveals a significant enrichment of 2 CREs showing significant homology with the binding site consensus of the transcription factor HY5 defined by DAP-seq (O'Malley et al., 2016) (Figure 7A, B). In total, 45 out of 108 genes involved in ROS detoxification under high N contained a putative HY5 binding site (Table S2 and S3). Interestingly, it has been shown previously that HY5 is involved in the control of ROS production in response to light or temperature treatments (Catala et al., 2011; Chen et al., 2013; Chai et al., 2015). To first validate the implication of HY5 in the regulation of genes involved in ROS detoxification under high N condition, we aimed to investigate the binding of HY5 to the putative consensus sequences identified in part of the ROS detoxification genes. To this end, we performed chromatin immunoprecipitation followed by quantitative PCR on a subset of genes with or without putative HY5 binding sites. The results showed that HY5 indeed binds to the promoter of detoxification genes containing a putative HY5 binding site, whereas no significant binding was observed for genes without putative HY5 binding site, although they are also involved in detoxification processes under high N provision (Figure 7C). Next, in order to test the effect of HY5 mutation, we measured the expression of these genes (with or without HY5 binding sites) in WT and in *hy5-215* mutant lines. Every gene we tested, except one, were statistically down-regulated in the *hy5-215* mutant line under high N provision (Figure 7D), strongly supporting a major role of HY5 in the activation of the detoxification program under high N provision. Finally, to test the effect of the mis-regulation of the detoxification program under high N following HY5 mutation, we measured the levels ^of ROS and H^2^O^2 ^in WT and in *hy5-215* mutant lines. The results clearly showed that ROS and H^2^O^2 ^levels were higher in *hy5-215* than in WT, demonstrating the functional importance of^ HY5 in the process of detoxification under high N provision (Figure 8). Altogether, our results demonstrate the induction of a transcriptional program needed for ROS detoxification under high N provision, which is driven in most part by the chromatin remodeler HNI9 and the transcription factor HY5.

**Figure 7:**
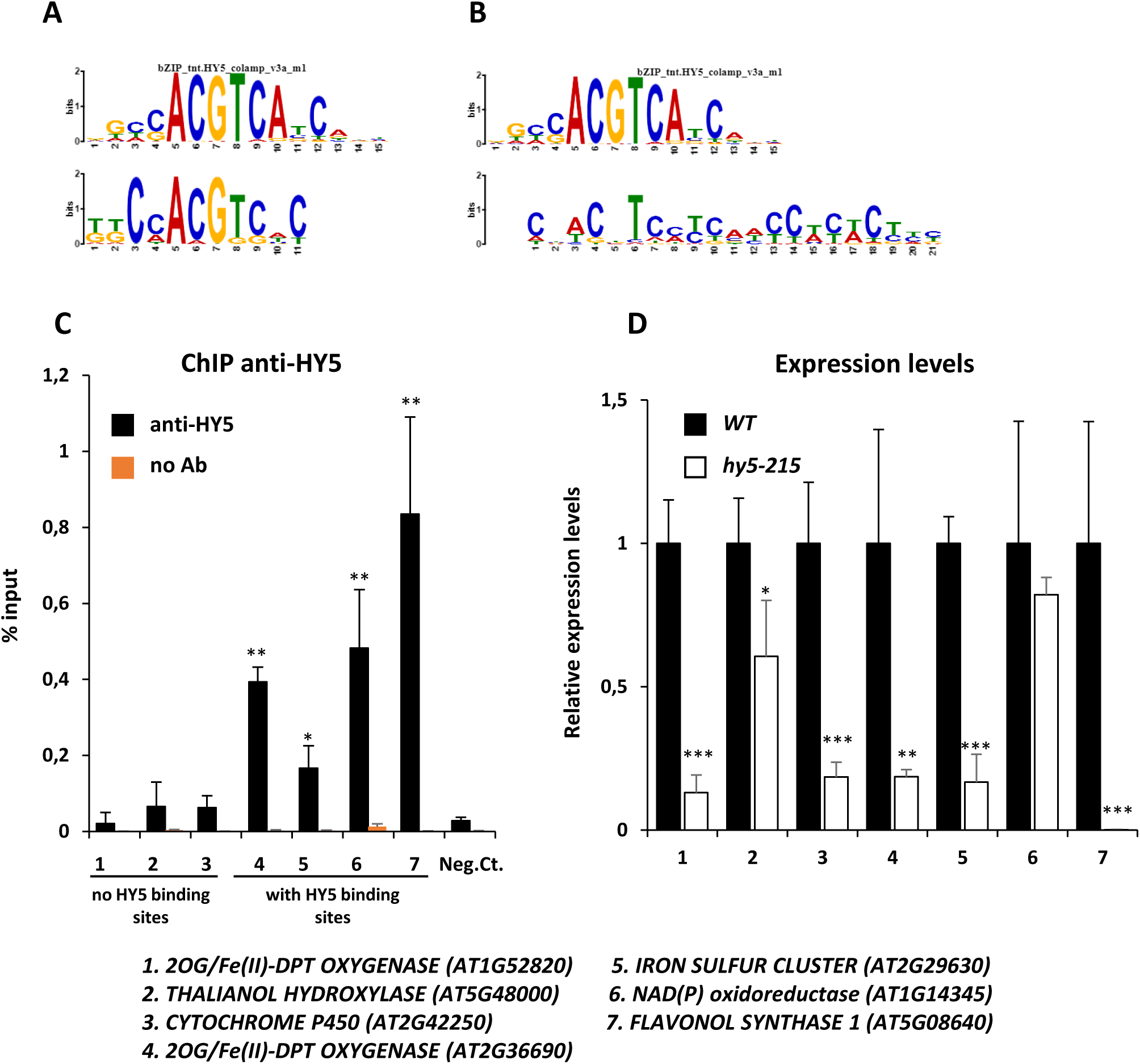
HY5 binds to and regulates the expression of genes involved in detoxification under high N provision. A, B. Comparison of 2 conserved motifs discovered by MEME analysis in the promoters of HNI9-dependent genes induced under high N provision (10 mM NH_4_NO_3_) with HY5 consensus binding site identified by DAP-seq. C. ChIP analysis of HY5 enrichment in WT roots at the loci of HNI9-dependent genes induced under high N provision. Quantification by qRT-PCR is shown as the percentage of input. Error bars represent standard errors of the mean based on at least 3 biological replicates. Statistical significance was computed using a two-tailed Student’s t-test (significance cutoff: *p < 0.05, **p < 0.01, ***p < 0.001), in comparison to a negative control (Neg. Ct.: *AT4G03900*, showing no relation with N or HY5 signaling). D. Relative expression of genes involved in detoxification induced under high N provision in roots from WT and *hy5-215* lines. Statistical significance was computed using a two-tailed Student’s t-test. Significance cutoff: *p < 0.05, **p < 0.01, ***p < 0.001.

**Figure 8:**
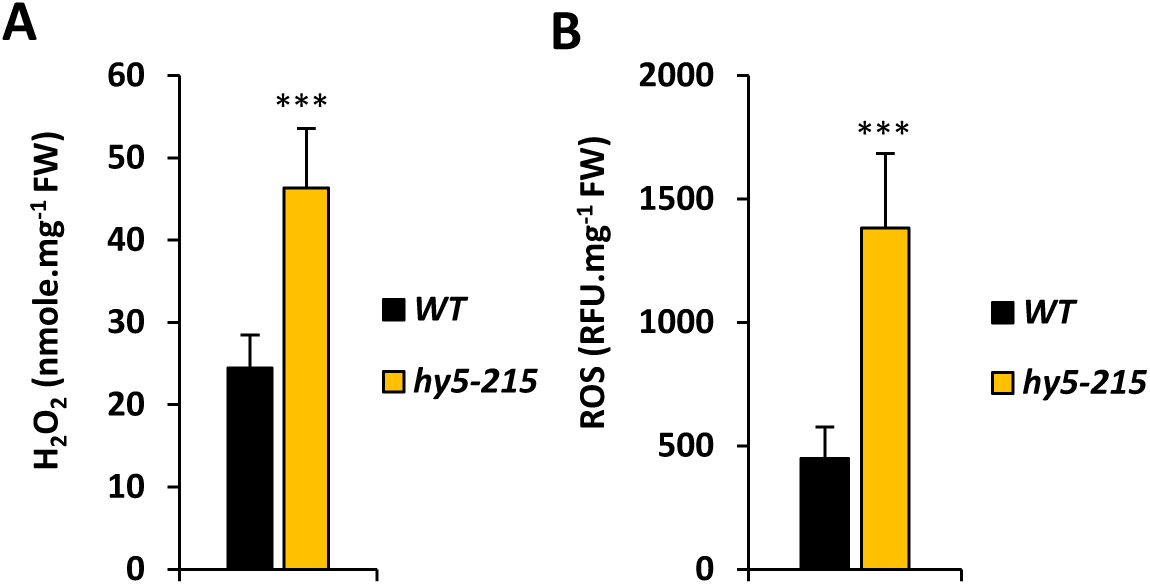
ROS levels are higher in *hy5-215* under high N provision. Measurement of H_2_O_2_(A) and ROS (B) in roots of WT and *hy5-215* lines under high N (10 mM NH_4_NO_3_) provision. Statistical significance was computed using a two-tailed Student’s t-test. Significance cutoff: *p < 0.05, **p < 0.01, ***p < 0.001.

## Discussion

Plants, like every living organism, have to cope with environmental constraints. ROS accumulation has been shown to be involved in numerous signaling pathways or to cause oxidative stress in response to different environmental challenges (Baxter et al., 2014; Choudhury et al., 2017; Noctor et al., 2018). The main finding of our work is that high N provision leads to the generation of ROS in plant roots, which are managed by the transcriptional induction a specific detoxification program. Interestingly, increasing N provision is viewed positively and as favorable conditions for plant growth, but we demonstrated here that excessive N nutrition could in fact be detrimental for the plant. Indeed, even if efficient detoxification systems exist and maintain ROS homeostasis, we can reasonably think that their induction and functioning imply a cost for plants, and that their efficiency can be limited under certain circumstances. In spite of the demonstration that high N nutrition leads to ROS accumulation, the physiological cause of ROS production under high N conditions remains to be determined. Nitrate assimilation and to a lower extent ammonium assimilation pathways can be candidate, as they consume a lot of reducing power (Hachiya and Sakakibara, 2017), and may in consequence alter the cellular redox balance of plants. However, it may also reflect a more general link between nutrition and redox balance. Indeed, several reports have described that excessive nutrition in animals leads to ROS production and perturbation of the cellular redox status (Sies et al., 2005; Samoylenko et al., 2013; Gorlach et al., 2015). In the case of plants, nutrient starvation has also been identified as a condition that generates oxidative stress through the production of ROS (Shin and Schachtman, 2004; Shin et al., 2005). In addition, recent reports demonstrated that ROS accumulate early during N starvation response, and most probably contribute to regulate nitrate-responsive genes like *NRT2.1* (Jung et al., 2018; Safi et al., 2018). Therefore, these studies and our present work suggest the existence of thresholds of N provision inside which the cellular redox balance is optimal. This situation is reminiscent of what has been described for iron (Fe) homeostasis. Indeed, excessive Fe provision can also lead to an accumulation of ROS, and specific mechanisms have been described to limit this negative effect (Briat et al., 2010). Nutrient deficiency or excess (at least for N and Fe) would correspond to conditions in which ROS production is above a physiological limit and alter plant redox status. Again, it is interesting to observe that the same analysis is true for animals (Gorlach et al., 2015).

In our work, we highlight the role of HNI9 in the transcriptional induction of this detoxification program. In Arabidopsis, HNI9 has been mainly associated to brassinosteroids (BRs) signaling (Li et al., 2010; Wang et al., 2014). In response to BRs, HNI9 is associated with the chromatin remodeling factors ELF6, REF6 and SDG8, and with the transcription factor BES1. ELF6 and REF6 are responsible for removing of the repressive chromatin mark H3K27me3, and SDG8 is supposed to catalyze the trimethylation of H3K36, which promotes elongation of transcription. This suggests a role for HNI9 in complexes associated with transcriptional switch, from repression to activation, as demonstrated in animals (Chen et al., 2012; Wang et al., 2013). In our work, HNI9 is mainly linked with differences in H3K4me3 levels. This suggests that HNI9 could be associated with other chromatin complexes or other steps of transcriptional regulation. Nevertheless, it identifies another transcriptional pathway in which HNI9 is essential for the induction of expression of a large set of genes.

In addition, our work identifies HY5 as a major component for the induction of the gene network involved in ROS detoxification under high N provision. HY5 is a master regulator of gene expression in Arabidopsis, at the center of several transcriptional networks. It is notably involved in the responses to photosynthesis, light, temperature or hormones (Gangappa and Botto, 2016). Interestingly, it has been also implicated in the control of ROS homeostasis in response to light or cold treatments (Catala et al., 2011; Chen et al., 2013; Chai et al., 2015). In each cases, HY5 is required to suppress ROS accumulation during stressful conditions. However, the set of genes induced by HY5 to balance ROS levels seems different from one stimulus to the others. This suggests that other transcription factors, in addition to HY5, could specify the response according to the environmental signal. Interestingly, we found that the expression of every gene of the ROS detoxification network in response to high N that we tested was dependent on HY5, whatever the presence of a HY5 binding site in their promoter. This may be due to the existence of several layers in this gene network, and suggests that HY5 would be at the top of the network.

In conclusion, we identified how a detoxification program is induced in order to maintain plant redox status under physiological conditions even under high nutritional provision. However, usage of detoxification processes is certainly not costless for plants. Thus, this work demonstrates that excessive N input is not appropriate for plant physiology and development, and lends support to the occurrence of an optimum between nutrition and ROS production in physiology.

## Material and methods

### Plant material and growth conditions

The *Arabidopsis thaliana* accession used in this study was Col-0. Mutant alleles and transgenic plants used in this study are *hni9-1* (Widiez et al., 2011), *vct2 (Collin et al., 2008)*, *hy5-215* (Oyama et al., 1997), and *GRX1-roGFP2* (Meyer et al., 2007). Experiments were performed using roots from 7 days-old seedlings grown under a long-day photoperiod (16 h light and 8 h dark) on vertical MS/2 plates without N (PlantMedia) supplied with 0.8 % agar, 0.1 % of sucrose, 0.5 g/L MES and the appropriate concentration of N as described in figures legend or in the main text.

### Analysis of gene expression by quantitative PCR

Root samples were frozen and ground in liquid nitrogen, and total RNA was extracted using TRI REAGENT (MRC), DNase treated (RQ1 Promega), and reverse transcription was achieved with M-MLV reverse transcriptase (RNase H minus, Point Mutant, Promega) using an anchored oligo(dT)20 primer. Transcript levels were measured by qRT-PCR (LightCycler 480, Roche Diagnostics) using the TB Green^TM^ Premix Ex Taq^TM^ (Tli Rnase H plus) Bulk (TaKaRa). Gene expression was normalized using *ACT2* as an internal standard. Sequences of primers used in qPCR for gene expression analysis are listed in Supplementary file 2.

### ChIP quantitative PCR

ChIP experiments were performed as previously described (Bellegarde et al., 2018). Chromatin was precipitated with 2.5 μg of antibodies against H3 (Abcam 1791), H3K27me3 (Millipore 07-449), H3K4me3 (Diagenode C15410030), H3K36me3 (Abcam 9050), H3K9ac (Agrisera AS163198), or HY5 (Agrisera AS12 1867). Immunoprecipitated DNA was purified by phenol-chloroform and ethanol-precipitated, and resulting DNA was analyzed by qPCR analysis. For chromatin marks analyses, ChIP experiments were quantified using H3 level as an internal standard, and normalized using *ACT7 (*H3K4me3, H3K9ac), *ACT2* (H3K36me3) or *LEC2* (H3K27me3) enrichment. For HY5 binding sites analyses, ChIP experiments were normalized to the input levels. Sequences of primers used in qPCR for ChIP experiments are listed in Supplementary file 2.

### ROS and H_2_O_2_assays

Root samples were frozen and ground in liquid nitrogen, and ROS and H_2_O_2_ were extracted using about 30 mg of plant material in 200 µL of phosphate buffer (20 mM K2HPO4, pH 6.5). For ROS measurements, 50 µL of supernatant were mixed with 50 µL of 20 µM DFFDA (Molecular Probes). Reactions were incubated 30 minutes at ambient temperature, and fluorescence was detected using 492/527 nm as excitation/emission parameters. H_2_O_2_ was measured using the Amplex Red Hydrogen Peroxide/Peroxidase Assay Kit (Invitrogen), as described in (Brumbarova et al., 2016).

### Microscopy

Confocal imaging of Arabidopsis root cells expressing *GRX1-roGFP2* were performed using a Leica SP8 Confocal Microscope (Leica, Germany). GFP quantifications were done with an Axiovert 200 M microscope (Zeiss, Germany) and images were analyzed with ImageJ analysis software. The data represent the fluorescence quantification values measured in the root tip.

### Root growth analyses

Vertical agar plates containing plants were scanned after 7 days of growth and root length was analyzed using ImageJ analysis software. For experiments with thiamin, plants were grown on vertical plates with MS/2 without N (PlantMedia) supplied with 0.8 % agar, 0.1 % of sucrose, 0.5 g/L MES and 10 mM NH_4_NO_3_ for 4 days, and transferred to the same medium with or without 100 µM thiamin. Position of the root tip was marked on the back of the new plates, and root length was analyzed 2 days after transferred by measuring growth from the mark to the root tip.

### Determination of conserved cis regulatory sequences

Putative cis regulatory sequences were identified using the MEME suite (Bailey et al., 2009). 500 base pairs of promoter sequences were used as primary input, with the following parameters: 0-order background model, classic discovery mode, 0 or one occurrence per sequence, motif width between 6 and 50 nucleotides.

### Analysis of gene ontology

Gene ontology analysis was performed using BINGO under Cytoscape environment, using Biological Process file, and a significance level of 0.05.

### Statistical analysis

Mean ± SE is shown for all numerical values. Statistical significance was computed using a two-tailed Student’s t-test. Significance cutoff: *p < 0.05, **p < 0.01, ***p < 0.001.

## Supporting information

## Acknowledgments

We thank members of A.G. lab for discussion. This work was supported by a grant from the National Agency for Research (ANR14-CE19-0008 *IMANA* to A.G., An.M., & G.K.). F.B. was the recipient of a PhD fellowship from INRA BAP department. We thank Andreas Meyer and Alexandre Martinière for the *GRX1-roGFP2* line, Fredy Barneche for the *hy5-215* line, and Jean-François Briat for the *vtc2* line. We thank the Montpellier Rio Imaging platform for microscopy observations.

