## Supplementary material for "HNI9 and HY5 maintain ROS homeostasis under high nitrogen provision in Arabidopsis"

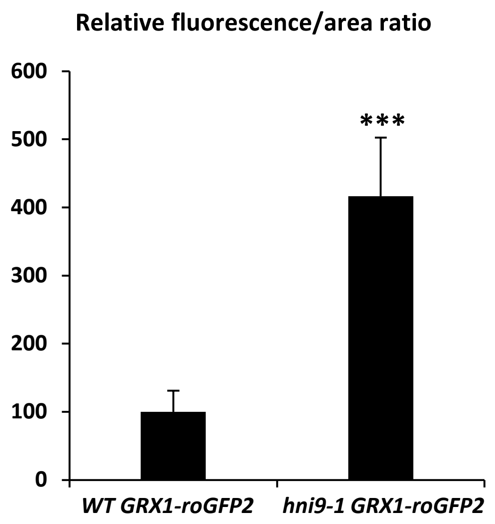


**Figure S1 : Quantification of GFP signal from *GRX1-roGFP2* probe in WT and *hni9-1* lines under high N (10 mM $NH_4NO_3$) provision**. Quantification of GFP signal was done using ImageJ software by measuring pixels values in the same area of root tips for each line. The data represent the mean pixel values of at least 15 plants. Statistical significance was computed using a two-tailed Student's t-test. Significance cutoff: *p < 0.05, **p < 0.01, ***p < 0.001.

| TAIR ID | TAIR Description |
|---|---|
| At5g48485 | DIR1, Bifunctional inhibitor/lipid-transfer protein/seed storage 2S albumin superfamily protein |
| At3g09220 | LAC7, laccase 7 |
| At1g51830 | Leucine-rich repeat protein kinase family protein |
| At3g62680 | ATPRP3, PRP3, proline-rich protein 3 |
| At5g47450 | ATTIP2;3, DELTA-TIP3, TIP2;3, tonoplast intrinsic protein 2;3 |
| At5g17230 | PSY, PHYTOENE SYNTHASE |
| At1g31710 | Copper amine oxidase family protein |
| At5g48000 | CYP708 A2, CYP708A2, THAH, THAH1, cytochrome P450, family 708, subfamily A, polypeptide 2 |
| At2g33830 | Dormancy/auxin associated family protein |
| At1g14345 | NAD(P)-linked oxidoreductase superfamily protein |
| At1g76790 | O-methyltransferase family protein |
| At1g21130 | O-methyltransferase family protein |
| At2g29630 | PY, THIC, thiaminC |
| At4g23670 | Polyketide cyclase/dehydrase and lipid transport superfamily protein |
| At5g48880 | KAT5, PKT1, PKT2, peroxisomal 3-keto-acyl-CoA thiolase 2 |
| At3g14990 | Class I glutamine amidotransferase-like superfamily protein |
| At2g36870 | XTH32, xyloglucan endotransglucosylase/hydrolase 32 |
| At2g42250 | CYP712A1, cytochrome P450, family 712, subfamily A, polypeptide 1 |
| At1g70890 | MLP43, MLP-like protein 43 |
| At2g43590 | Chitinase family protein |
| At2g39040 | Peroxidase superfamily protein |
| At1g74670 | Gibberellin-regulated family protein |
| At1g12220 | RPS5, Disease resistance protein (CC-NBS-LRR class) family |
| At3g49780 | ATPSK3 (FORMER SYMBOL), ATPSK4, PSK4, phytosulfokine 4 precursor |
| At2g03470 | ELM2 domain-containing protein |
| At4g14060 | Polyketide cyclase/dehydrase and lipid transport superfamily protein |
| At2g33850 | unknown protein; FUNCTIONS IN: molecular_function unknown; INVOLVED IN: biological_process unknown; LOCATED IN: endomembrane system; EXPRESSED IN: 17 plant structures; EXPRESSED DURING: 9 growth stages; BEST Arabidopsis thaliana protein match is: unknown protein (TAIR:AT1G28400.1); Has 3053 Blast hits to 2119 proteins in 133 species: Archae - 6; Bacteria - 52; Metazoa - 135; Fungi - 96; Plants - 73; Viruses - 2; Other Eukaryotes - 2689 (source: NCBI BLink). |

| | | |
|---|---|---|
| | At5g62165 | AGL42, AGAMOUS-like 42 |
| | At5g60100 | APRR3, PRR3, pseudo-response regulator 3 |
| | At5g50760 | SAUR-like auxin-responsive protein family |
| | At3g43960 | Cysteine proteinases superfamily protein |
| | At5g57220 | CYP81F2, cytochrome P450, family 81, subfamily F, polypeptide 2 |
| | At4g22610 | Bifunctional inhibitor/lipid-transfer protein/seed storage 2S albumin superfamily protein |
| | At3g12955 | SAUR-like auxin-responsive protein family |
| | At2g24160 | pseudogene, leucine rich repeat protein family, contains leucine rich-repeat domains Pfam:PF00560, INTERPRO:IPR001611; contains some similarity to Cf-4 (Lycopersicon hirsutum) gi\|2808683\|emb\|CAA05268; blastp match of 37% identity and 8.4e-98 P-value to GP\|2808683\|emb\|CAA05268.1\|\|AJ002235 Cf-4 {Lycopersicon hirsutum} |
| | At1g14240 | GDA1/CD39 nucleoside phosphatase family protein |
| | At4g23680 | Polyketide cyclase/dehydrase and lipid transport superfamily protein |
| | At4g00780 | TRAF-like family protein |
| | At1g23030 | ARM repeat superfamily protein |
| | At5g08640 | ATFLS1, FLS, FLS1, flavonol synthase 1 |
| | At3g16470 | JR1, Mannose-binding lectin superfamily protein |
| | At1g73780 | Bifunctional inhibitor/lipid-transfer protein/seed storage 2S albumin superfamily protein |
| | At1g80520 | Sterile alpha motif (SAM) domain-containing protein |
| | At2g34490 | CYP710A2, cytochrome P450, family 710, subfamily A, polypeptide 2 |
| | At2g41480 | Peroxidase superfamily protein |
| | At4g23496 | SP1L5, SPIRAL1-like5 |
| | At5g30440 | transposable element gene |
| | At5g42900 | COR27, cold regulated gene 27 |
| | At1g13930 | Involved in response to salt stress. Knockout mutants are hypersensitive to salt stress. |
| | At5g57630 | CIPK21, SnRK3.4, CBL-interacting protein kinase 21 |
| | At3g13950 | unknown protein; BEST Arabidopsis thaliana protein match is: unknown protein (TAIR:AT4G13266.1); Has 339 Blast hits to 265 proteins in 12 species: Archae - 0; Bacteria - 0; Metazoa - 0; Fungi - 0; Plants - 339; Viruses - 0; Other Eukaryotes - 0 (source: NCBI BLink). |
| | At1g19960 | BEST Arabidopsis thaliana protein match is: transmembrane receptors (TAIR:AT2G32140.1); Has 41 Blast hits to 41 proteins in 17 species: Archae - 0; Bacteria - 2; Metazoa - 23; Fungi - 0; Plants - 11; Viruses - 0; Other Eukaryotes - 5 (source: NCBI BLink). |
| | At3g48350 | Cysteine proteinases superfamily protein |
| | At3g60280 | UCC3, uclacyanin 3 |

| | |
|---|---|
| At1g62480 | Vacuolar calcium-binding protein-related |
| At1g75780 | TUB1, tubulin beta-1 chain |
| At1g20620 | ATCAT3, CAT3, SEN2, catalase 3 |
| At2g37130 | Peroxidase superfamily protein |
| At2g47600 | ATMHX, ATMHX1, MHX, MHX1, magnesium/proton exchanger |
| At5g19870 | Family of unknown function (DUF716) |
| At1g60470 | AtGolS4, GolS4, galactinol synthase 4 |
| At2g45920 | U-box domain-containing protein |
| At5g24660 | LSU2, response to low sulfur 2 |
| At1g15170 | MATE efflux family protein |
| At1g72180 | Leucine-rich receptor-like protein kinase family protein |
| At1g12090 | ELP, extensin-like protein |
| At1g52820 | 2-oxoglutarate (2OG) and Fe(II)-dependent oxygenase superfamily protein |
| At1g51840 | protein kinase-related |
| At4g30670 | Putative membrane lipoprotein |
| At3g23400 | FIB4, Plastid-lipid associated protein PAP / fibrillin family protein |
| At2g23910 | NAD(P)-binding Rossmann-fold superfamily protein |
| At3g21560 | UGT84A2, UDP-Glycosyltransferase superfamily protein |
| At5g42600 | MRN1, marneral synthase |
| At4g13420 | ATHAK5, HAK5, high affinity K+ transporter 5 |
| At5g56870 | BGAL4, beta-galactosidase 4 |
| At4g03330 | ATSYP123, SYP123, syntaxin of plants 123 |
| At4g29270 | HAD superfamily, subfamily IIIB acid phosphatase |
| At1g54970 | ATPRP1, PRP1, RHS7, proline-rich protein 1 |
| At5g47000 | Peroxidase superfamily protein |
| At1g52410 | TSA1, TSK-associating protein 1 |
| At2g16750 | Protein kinase protein with adenine nucleotide alpha hydrolases-like domain |
| At4g15480 | UGT84A1, UDP-Glycosyltransferase superfamily protein |
| At1g75280 | NmrA-like negative transcriptional regulator family protein |
| At4g16260 | Glycosyl hydrolase superfamily protein |
| At4g17340 | DELTA-TIP2, TIP2;2, tonoplast intrinsic protein 2;2 |
| At2g16005 | MD-2-related lipid recognition domain-containing protein |
| At1g68500 | unknown protein; BEST Arabidopsis thaliana protein match is: unknown protein (TAIR:AT1G25422.1); Has 16 Blast hits to 16 proteins in 6 species: Archae - 0; Bacteria - 0; Metazoa - 0; Fungi - 0; Plants - 16; Viruses - 0; Other Eukaryotes - 0 (source: NCBI BLink). |
| At1g10090 | Early-responsive to dehydration stress protein (ERD4) |

| | |
|---|---|
| At5g54510 | DFL1, GH3.6, Auxin-responsive GH3 family protein |
| At3g16240 | AQP1, ATTIP2;1, DELTA-TIP, DELTA-TIP1, TIP2;1, delta tonoplast integral protein |
| At3g29780 | RALFL27, ralf-like 27 |
| At4g11360 | RHA1B, RING-H2 finger A1B |
| At3g60190 | ADL1E, ADL4, ADLP2, DL1E, DRP1E, EDR3, DYNAMIN-like 1E |
| At3g27200 | Cupredoxin superfamily protein |
| At1g18870 | ATICS2, ICS2, isochorismate synthase 2 |
| At4g37410 | CYP81F4, cytochrome P450, family 81, subfamily F, polypeptide 4 |
| At2g41970 | Protein kinase superfamily protein |
| At4g20320 | CTP synthase family protein |
| At4g23700 | ATCHX17, CHX17, cation/H+ exchanger 17 |
| At5g17040 | UDP-Glycosyltransferase superfamily protein |
| At1g21100 | O-methyltransferase family protein |
| At1g64370 | unknown protein; Has 773 Blast hits to 375 proteins in 118 species: Archae - 0; Bacteria - 97; Metazoa - 421; Fungi - 108; Plants - 31; Viruses - 0; Other Eukaryotes - 116 (source: NCBI BLink). |
| At1g05200 | ATGLR3.4, GLR3.4, GLUR3, glutamate receptor 3.4 |
| At3g15090 | GroES-like zinc-binding alcohol dehydrogenase family protein |
| At2g36690 | 2-oxoglutarate (2OG) and Fe(II)-dependent oxygenase superfamily protein |
| At5g19860 | Protein of unknown function, DUF538 |
| At1g11970 | Ubiquitin-like superfamily protein |
| At5g48485 | DIR1, Bifunctional inhibitor/lipid-transfer protein/seed storage 2S albumin superfamily protein |

**Table S1**: List of 108 genes induced under high N condition (10 mM $NH_4NO_3$) in a HNI9-dependent manner, and the corresponding description of their function in TAIR.

| Name | Strand | Start | *p*-value | Sites | | |
|---|---|---|---|---|---|---|
| 2. AT1G10090 | + | 431 | 5.68e-8 | ATTCACGTTA | TGCCACGTCAC | ATGATCTGGC |
| 90. AT5G19860 | - | 408 | 8.42e-8 | TAGACGTAAT | TGCCACGTCTC | ATCTAAAATT |
| 106. AT5G60100 | + | 376 | 1.12e-7 | TTTTGATTTT | GTCCACGTCAC | TAATCTCAGC |
| 100. AT5G48880 | + | 476 | 7.28e-7 | AATTGCCAAA | TGCAACGTCAC | TTCTTTCACT |
| 9. AT1G15170 | - | 45 | 7.83e-7 | TATCCTGATC | GTCCACGTGTC | AAAACATATG |
| 5. AT1G12220 | + | 127 | 9.96e-7 | TTTTCTATTA | TTCCACGGCCC | AACTTCGGTT |
| 8. AT1G14345 | + | 433 | 1.07e-6 | ATAGAGCTTA | GGCCACGTGGC | ACAATTTGCG |
| 71. AT4G03330 | + | 416 | 1.60e-6 | CATTATTGGA | GACCACGTCTC | TGCAACATGG |
| 43. AT2G36690 | + | 446 | 1.75e-6 | GACTATTCAT | TTCAACGTCTC | CTCCAAAAAC |
| 95. AT5G42900 | - | 445 | 1.92e-6 | TGGGTAAAGT | TGCCACGTCAT | CACCAGTCAA |
| 60. AT3G21560 | + | 281 | 2.11e-6 | AAATTTCTGT | ATCCACGTCAC | CTACTGACCC |
| 54. AT3G12955 | - | 59 | 2.63e-6 | ACATAAAGTA | TGCAACGGCTC | AAGCGAAGCT |
| 56. AT3G14990 | - | 343 | 3.34e-6 | TTTTGAGTCT | TTCCACGTCAT | TGTATTGGAT |
| 36. AT2G16750 | - | 367 | 3.60e-6 | GTTTTAAAAG | GGCCAAGTCCC | AAGTGAATAT |
| 7. AT1G14240 | + | 346 | 3.60e-6 | AACCACTCAA | GTCTACGTCAC | AAAAAAAATC |
| 55. AT3G13950 | - | 107 | 5.98e-6 | TCACTCTTCT | TTCAACGTGTC | TTTTCTTTCA |

**Table S2**: Genes from the list of 108 genes induced under high N condition (10 mM $NH_4NO_3$) in a HNI9-dependent manner showing the presence of a putative HY5 binding site (motif #1) in their promoter. Genes tested by ChIP-qPCR are underlined.

| Name | Strand | Start | *p*-value | Sites | | |
|---|---|---|---|---|---|---|
| 25. AT1G68500 | + | 451 | 4.76e-11 | CTTCCACTAA | CTACTTCCTCAACCTCTCCCT | TATAAATAAA |
| 37. AT2G23910 | + | 465 | 7.62e-9 | CTATTTATTA | CTACTTCCTTCTCCTCACGTC | TCTCCTCTAC |
| 3. AT1G11970 | - | 80 | 8.66e-9 | AAAAGTTCTT | CATCTTCCTCGTCCTCACTTT | ATTCCTATTC |
| 68. AT3G60280 | + | 434 | 1.43e-8 | AGTCTGTAGT | CGTCATCGTCCTCCTCACCCC | ATAGCATCTC |
| 48. AT2G41970 | - | 362 | 1.43e-8 | ACGTCTTGCA | CGACCTCTTCAACCATACTTC | TAATTTATCT |
| 87. AT5G08640 | + | 452 | 3.70e-8 | CAAGATTTCG | CCACGTCCTCACTTCCTCCCT | CCTTAAAACC |
| 75. AT4G15480 | - | 374 | 5.83e-8 | TATTCAACAC | CTACTTGCTCACTCTCTCTCG | ACCACACATC |
| 58. AT3G16240 | + | 440 | 5.83e-8 | AATTCGGGTC | CAACGTAACCCACCACACCAC | AGAAACTCAT |
| 1. AT1G05200 | + | 417 | 1.39e-7 | CCTTGGCCTA | CGTCTTCACTGACCACTCCAG | AAATAGCACT |
| 37. AT2G23910 | + | 435 | 1.55e-7 | CAACAATGAA | ATAGTTCACCCACCTTTCTCC | TATTTATTAC |
| 100. AT5G48880 | - | 426 | 3.85e-7 | GCCATACTCT | TTTGATCCTCCACCACTCTTC | CTACCAACCT |
| 72. AT4G11360 | + | 35 | 4.25e-7 | ACCATCTGAG | CTACATCCCCATTTGCTCTTC | TATATGAAAT |
| 30. AT1G75280 | - | 248 | 4.69e-7 | TCGATTGACC | CCAGGTCCCCAACCCTTAGCC | CCCAACAGGA |
| 39. AT2G29630 | + | 410 | 5.16e-7 | ACTCGTGAAA | CGACGTTCTCCTCCTCACGTA | CCTTATCTTA |
| 23. AT1G62480 | + | 79 | 5.16e-7 | CCATCGATTT | TCTCCTCACCATCATCACTCC | TTCAAATATT |
| 67. AT3G60190 | + | 139 | 5.68e-7 | AGCCTCCTAC | CCTCTTCCTGGACCTAACTTT | ATTATTGGGT |
| 41. AT2G33850 | + | 411 | 8.29e-7 | GTCCTTATTT | CCACCTAACCATTTACACTCT | ATATAGTAAG |
| 40. AT2G33830 | + | 30 | 9.10e-7 | TTCCACACTT | CTCCCTCACTGCTCACGCTTC | TCATGATCAA |
| 68. AT3G60280 | + | 383 | 1.09e-6 | ATCAAAGCCA | CGACTTCTTCAAACATTCTTC | TTTACTAACT |
| 11. AT1G19960 | - | 102 | 1.44e-6 | TCAACCTACA | CCACTCACTGACTCACACCAC | CATTATTGAG |
| 3. AT1G11970 | - | 179 | 1.44e-6 | CGCAGACCAA | CGTCACCCTCAACCTTTTTCT | ATTCACCCAA |
| 50. AT2G43590 | + | 472 | 1.87e-6 | ACCCCTATAT | ATACCTCACCACCTTTGCCCT | CTCAACCA |
| 26. AT1G70890 | + | 64 | 2.05e-6 | TCGTGCCTTG | ATTCCTCCTCTACCACTTTCG | GTGGGCTTTA |
| 26. AT1G70890 | + | 446 | 2.23e-6 | CTATCATTTA | ATACTTTCTCACTCATTCTTC | TACAAATAGA |
| 72. AT4G11360 | + | 162 | 2.43e-6 | ATCGTGTTCA | TATCTTCACTATCCACTCTTG | AATCCCTGAT |
| 100. AT5G48880 | + | 478 | 2.65e-6 | TTGCCAAATG | CAACGTCACTTCTTTCACTCC | TC |
| 88. AT5G17040 | + | 447 | 2.65e-6 | ATTTAATCCA | CAAGGTCCACAACCATTCCAT | ATTCATATTC |
| 15. AT1G23030 | - | 402 | 2.65e-6 | TTTTAAGGAC | CTACGTCTTGGTCCACTTTTT | TTGTTGTTGT |
| 106. AT5G60100 | + | 378 | 2.89e-6 | TTGATTTTGT | CCACGTCACTAATCTCAGCAT | GCGTAAATTT |
| 41. AT2G33850 | + | 180 | 3.15e-6 | CTAATATATA | CTACAACCCCATCCTTACGGG | GCATGGTGCT |
| 9. AT1G15170 | - | 278 | 3.15e-6 | TGTATTTTGA | CCTTTTTATCATCCACACTCT | TTTTCAATCT |
| 30. AT1G75280 | - | 419 | 3.42e-6 | TTACGATAAT | ACCCTTACTGAACCTTACTTC | AATTCTAAGA |

**Table S3:** Genes from the list of 108 genes induced under high N condition (10 mM $NH_4NO_3$) in a HNI9-dependent manner showing the presence of a putative HY5 binding site (motif #2) in their promoter. Genes tested by ChIP-qPCR are underlined.
